# Long-range regulatory sequence models reveal a phylogenetic boundary in vertebrate genome evolution

**DOI:** 10.64898/2026.11.09.765190

**Authors:** Binh Le, Hoang Nguyen, Lars Grønvold, Simen Rød Sandve, Oleksandr Frei, Ole A. Andreassen, Tuyen Trung Truong, Dat Thanh Nguyen

## Abstract

Regulatory activity at a locus depends on local sequence and on long-range interactions with sequence further away, but how long each kind of information stays shared between genomes over evolution is unknown. Here we use sequence-to-function models as a comparative assay, training FunFormer, a long-context binary classifier that reads 20 kb, together with an architecture- and parameter-matched control that reads only the central 1 kb, in six vertebrates spanning 435 million years, and measuring what each retains when applied to another genome. Within species, the long-context model outperformed three published 1 kb classifiers by 1.0 to 3.8 percentage points in every genome, a gain that came from context rather than architecture. Across species, its contribution decayed with divergence, was near zero at the amniote split and reversed between 320 and 435 Mya, while local sequence still transferred into fish. Zebrafish and Atlantic salmon, which share their divergence from tetrapods but differ by the salmonid genome duplication, behaved alike, so the reversal is not explained by the salmonid duplication. An *in silico* flank intervention showed that replacing the flank with GC-matched sequence from the target genome cost 8 percentage points among mammals. When a mammalian model was applied to fish, a flank drawn from the source genome on average performed as well as the target’s own flank, so the model drew nothing from the fish flank that an arbitrary source-genome flank did not provide. The long-range information that these models learn therefore stops being shared between vertebrate genomes well before the local information does. The same boundary sets a practical rule for reusing models in species without their own data, where long context helps among close relatives but is a liability across deep splits, and in variant scoring most foreign models matched a native model in data-limited pig but none did in human.

## 1 Introduction

Regulatory activity at a locus is encoded both in local sequence at the site, which carries the transcription factor binding sites that drive its activity [1], and in sequence further away that contributes through long-range interactions [2]. Models that read more of the genome therefore predict regulatory activity more accurately [3, 4], but the two scales need not be conserved for equally long. Transcription factors and their binding preferences are deeply conserved across vertebrates, yet regulatory elements turn over rapidly, and only about 1% of human liver enhancers retained activity across ten placental mammals [5]. In salmonids, duplicated enhancers also diverged faster than duplicated promoters after the most recent whole-genome duplication [6]. At the same time, long-range regulatory domains around some developmental genes have kept their synteny from teleosts to mammals [7], but these observations concern particular classes of elements and loci. How long the long-range regulatory information around a typical locus stays shared between genomes, and whether it outlasts the local information or is lost before it, has not, to our knowledge, been measured genome-wide.

Sequence-to-function models offer a way to measure this, because what a model can learn depends on how much sequence it sees. They predict regulatory activity directly from DNA and have become central tools for interpreting the noncoding genome and for variant effect prediction (VEP) [4, 8, 9]. Classification models such as DeepSEA [8] and DanQ [10] predict whether a locus overlaps a peak of transcription factor binding, chromatin accessibility or histone modification from roughly one kilobase (kb) of local sequence. Regression models extended the window to capture long-range interactions, from Basenji at 131 kb [3] to Enformer at about 200 kb [4], Borzoi at about 500 kb [11] and AlphaGenome at one megabase [9]. The gains of Enformer over its short-range predecessors were attributed to integrating exactly those interactions [4]. A model given a long window can use long-range information, whereas a model given 1 kb can use only local sequence. If both models are trained in one species and applied to the genome of another, the difference in their accuracy there is the value of long-range information in the target, positive if what the long-context model learned still applies and negative if it no longer does. The short-context model alone shows how well local information transfers, and repeating this for every pair of species turns cross-species transfer into a comparative measurement of how long each kind of regulatory information stays shared.

Cross-species transfer has so far been studied as a question of model reuse, and within mammals it works well. Joint training on human and mouse improves prediction in both [12], enhancer classifiers trained in one mammal predict enhancers in another almost as well as within-species models [13], and enhancer logic generalizes where sequence alignment fails [14]. Models of tissue-specific open chromatin trained on a few mammals have been applied to predict regulatory conservation across more than 200 mammals [15], deep learning on matched functional profiles maps functional conservation between human and pig [16], and human-trained accessibility models predict accessibility in other primates almost as well as models trained in those species and have been used to find regions with increased accessibility in humans [17]. Beyond Mammalia, human- and mouse-trained models annotate chromatin in cattle, pig and chicken far better than in European seabass, with H3K4me3 transferring best [18], and a Sei-based model in plants shows the same decline with distance [19], while a plant DNA language model trained across 65 angiosperm genomes gains from longer context in large genomes [20]. A deep-learning model of chromatin accessibility in five vertebrates concludes that regulatory grammar is more conserved than the sequence carrying it [21], and mammalian weights help initialize a zebrafish expression model [22]. These studies establish that absolute accuracy declines with distance, but they cannot be read as a measurement of how long each kind of regulatory information stays shared, for two reasons. Where the models compared differ in context, they also differ in architecture, output type and training data, as when an Enformer trained jointly on human and mouse is set against a 1 kb DeepSEA, and joint human-mouse training alone improves held-out accuracy in both species [12]. Absolute accuracy in a target genome also confounds how transferable the source model is with how predictable the target is at all, which only a model trained on the target itself can separate. The question has practical stakes as well, because most species lack the epigenomic compendia needed to train a model *de novo*, so reusing a model from a data-rich species is an attractive route to genome interpretation in livestock, aquaculture species and other non-model organisms.

Such a measurement needs a binary model operating at long range. Continuous predictions are hard to compare across species because sequencing depth, normalization and dynamic range differ between datasets and genomes, so the outputs of a regression model have no shared scale. Regression models of liver accessibility across five mammals generalize to held-out species in aggregate yet predict quantitative differences at orthologous regions poorly, and the cross-species tasks that have succeeded so far have been binary [23]. Whether a locus is active is a label defined within each species, and a ranking metric over matched regulatory channels compares models without any shared scale. A long-context binary classifier, paired with a short-context control of identical architecture, is therefore the instrument the question needs.

Here we introduce FunFormer, a long-context binary classifier whose convolutional encoder compresses up to 50 kb of sequence into a fixed set of tokens for transformer layers with rotary positional encodings and bidirectional attention. Because the number of tokens is fixed, its 1 kb and 20 kb configurations have identical parameter counts, so FunFormer-1k is an architecture- and parameter-matched control and the difference between the two isolates context length. The fixed token count is also what makes long context practical for a binary classifier, since context can grow from 1 to 50 kb at constant parameter count. We trained both, together with three published 1 kb baselines, in six vertebrates spanning about 435 million years, namely human, pig, cattle, chicken, Atlantic salmon and zebrafish (Fig. 1). Pairs are grouped by the node at which they diverge, as mammal-mammal (58 Mya for pig-cattle, 94 Mya for human against either), mammal-chicken (320 Mya, the amniote split) and tetrapod-fish (435 Mya). The groups are nested rather than disjoint, since mammals are tetrapods too and chicken is a tetrapod but not a mammal. The deepest is named for the clade wide enough to cover every pair in it, chicken-fish included. The two fish are a matched pair, both teleosts at the same 435 Mya split from every tetrapod here and both carrying the teleost genome duplication (Ts3R) [24], but only salmon carrying the later salmonid round (Ss4R) [25], so comparing transfer into the two separates evolutionary distance from the salmonid duplication. For every ordered species pair we apply a model trained in one species to the genome of another and score matched (assay, tissue) channels by the area under the receiver operating characteristic curve (AUROC).

**Figure 1:**
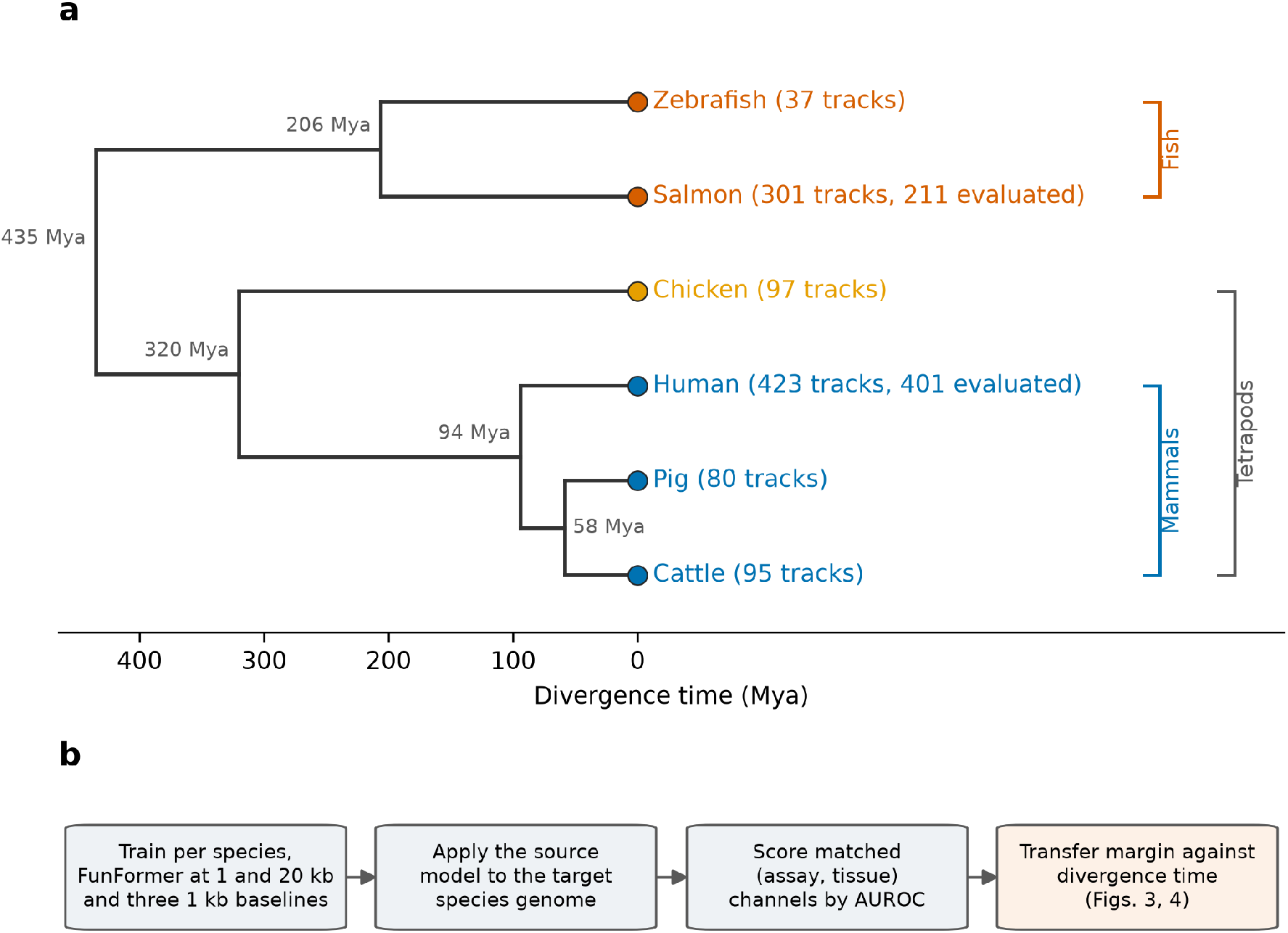
Study design. **(a)** Time-calibrated phylogeny of the six vertebrates, node ages in millions of years (Mya), tips colored by clade, with tracks trained on and, where fewer, tracks evaluated. Brackets mark clades (fish, mammals and tetrapods), mammals nesting inside tetrapods. Both fish sit at the same 435 Mya split from every tetrapod and share the teleost genome duplication, but only salmon carries the salmonid round. **(b)** Evaluation pipeline. Within each species we train FunFormer at 1 and 20 kb and the three published 1 kb baselines, then apply each to a target genome, score matched (assay, tissue) channels by AUROC and summarize as a transfer margin against divergence time.

We find that FunFormer outperforms three published 1 kb classifiers within every species, a gain that comes from context rather than architecture, but that across species the value of that context decays with divergence, is near zero at the amniote split and reverses before the tetrapod-fish split, while the 1 kb model still transfers into fish. Zebrafish and salmon behave alike, so the reversal is not explained by the salmonid duplication. An *in silico* flank intervention shows what the long context carries. Among mammals the model reads locus-specific information in the flank, whereas when a mammalian model is applied to a fish genome it draws on average no more from the target’s own flank than from an arbitrary stretch of the genome it was trained on. The long-range information that these models learn therefore stops being shared between vertebrate genomes well before the local information does. The same boundary sets a practical rule for model reuse, and in a downstream task, molecular QTL prioritization, a foreign model scores variants in data-limited pig about as well as a pig-trained model but falls clearly short in human.

## 2 Results

### 2.1 Overview of the study

We developed FunFormer, a deep neural network that predicts binary regulatory-element activity directly from DNA sequence while efficiently incorporating long-range sequence context (Figure 1), and we use it to address three aims. First, we build a long-context model for binary regulatory-element prediction and ask whether it improves on established architectures. Second, we evaluate how well such models transfer across species, what that transfer reveals about the evolutionary conservation of regulatory information, and, by ablating the sequence outside the central 1 kb, what the long context is being used for. Third, we assess zero-shot VEP, asking how well a sequence-to-function model trained only on regulatory activity prioritizes functional variants without ever seeing variant labels, and how this depends on the genome it was trained in.

We benchmarked FunFormer against DeepSEA [8], DeepFormer [26], and DeepATT [27], three widely used sequence-to-function architectures, and against FunFormer-1k, the same FunFormer architecture run at the baselines’ own 1 kb context. Because Fun-Former compresses any input length to a fixed number of tokens, its 1 kb and 20 kb configurations have identical parameter counts. FunFormer-1k is therefore an architecture- and parameter-matched control that separates the effect of context length from the effect of architecture throughout this study. We assembled epigenomic data for six vertebrates spanning roughly 435 million years of evolution, namely human, pig, cattle, chicken, salmon, and zebrafish (Figure 1a). Each dataset comprises 37 to 423 binary regulatory tracks covering chromatin accessibility and histone modifications. Human tracks are from the EN-TEx resource [28], the cattle, pig, chicken and salmon tracks are the FAANG- and AQUA-FAANG-derived processed datasets described in our earlier work [29], and the zebrafish tracks are from a published map of zebrafish cis-regulatory elements [30]. We trained all five models within each species, then transferred each source model onto the genomes of the other species and scored performance on matched (assay, tissue) channels using the area under the receiver operating characteristic curve (AUROC; Figure 1b). Finally, we tested zero-shot VEP in cross-species settings on putative causal expression QTL (eQTL) variants in human and pig and splicing QTL (sQTL) variants in pig, each paired with a matched control set. For every within- and cross-species model, we used the predicted change in regulatory activity as a feature to classify causal variants from controls. This asks whether the phylogenetic pattern seen in regulatory-element transfer carries over to variant-level prediction.

### 2.2 FunFormer enhances regulatory-element prediction within species

A model is a credible instrument for cross-species comparison only if it is a strong predictor within each genome. We therefore first evaluated within-species accuracy on held-out test intervals in all six species (Figure 2a,b). FunFormer-20k achieved the highest mean AUROC of the five models in every species, with 0.967 in human, 0.955 in salmon, 0.950 in zebrafish, 0.949 in cattle, 0.927 in pig, and 0.876 in chicken (Supplementary Table S1), exceeding the best published 1 kb baseline by 1.0 to 3.8 percentage points (pp). This advantage could come from the transformer architecture, from the longer context, or both. FunFormer-1k, the same architecture run at the baselines’ own 1 kb context, separates the two. Averaged over the six species, replacing the best published baseline with FunFormer-1k changed mean AUROC by +0.19 pp (range *−* 0.01 to +0.42 pp), whereas extending FunFormer-1k to 20 kb added +1.8 pp (range +0.8 to +3.6 pp). At matched context FunFormer is no better than existing 1 kb models. Essentially all of its gain comes from context length. FunFormer-1k is therefore the 1 kb reference for every cross-species comparison later in the text, with the best published baseline reported alongside.

**Figure 2:**
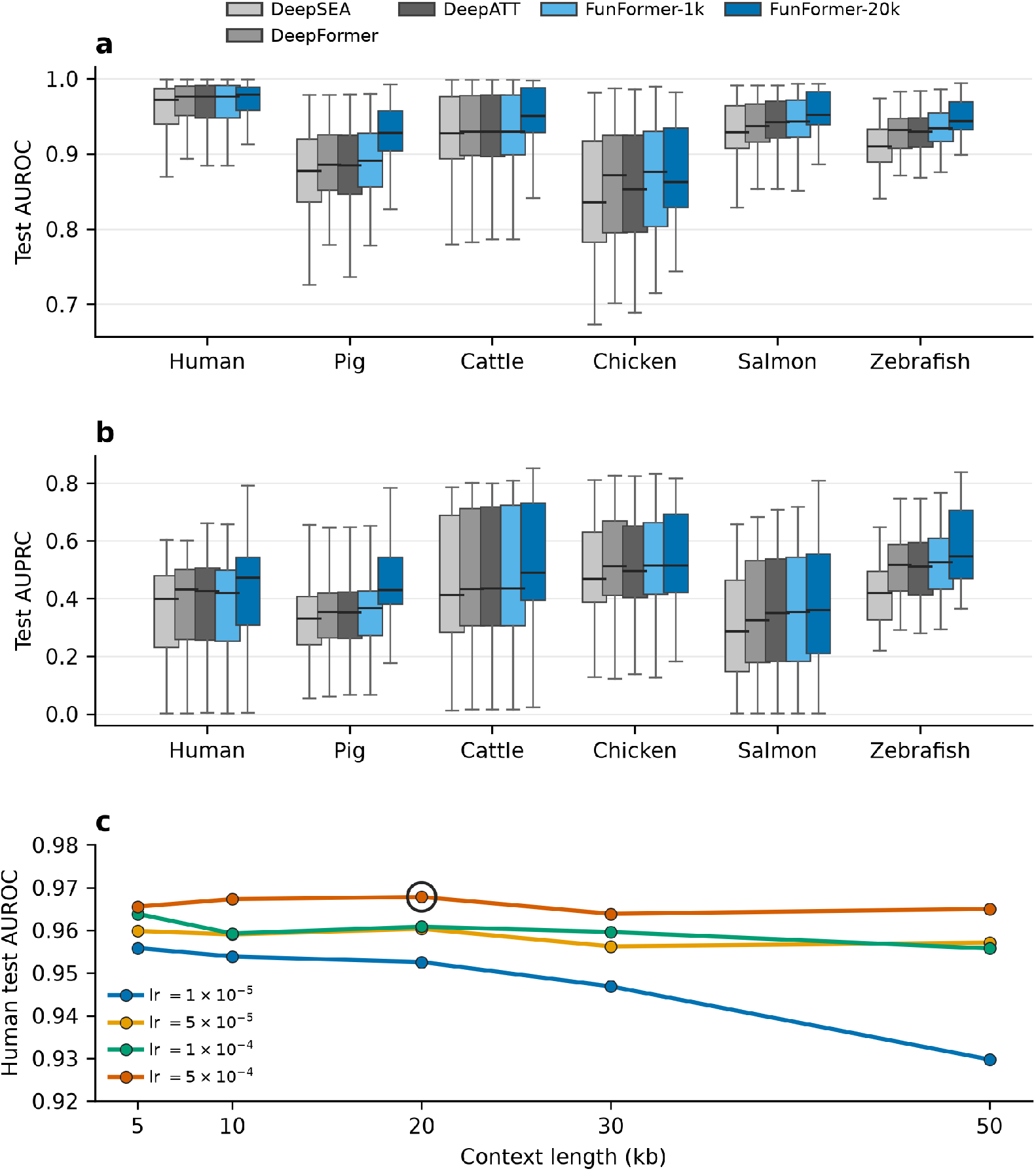
FunFormer enhances regulatory-element prediction within species. **(a**,**b)** Per-track test AUROC (a) and AUPRC (b) for the five models in each species (box, interquartile range; line, median; whiskers, 1.5*×* that range). FunFormer-20k has the highest mean AUROC and the highest median AUPRC in every species. In chicken, FunFormer-1k leads on mean AUPRC and median AU-ROC. **(c)** Human context-length ablation, 5 to 50 kb by four learning rates, 20 runs. The circled point is the selected configuration, 20 kb at 5 *×* 10^*−*4^. FunFormer-1k was trained separately and is not in this sweep. The sweep is scored on every track, without the two filters described in Methods that panels a and b and Supplementary Table S1 apply. Its values are therefore comparable with one another but sit slightly above the table, at 0.968 for the selected run against 0.967 there.

The advantage held for the area under the precision-recall curve (AUPRC), which is dominated by the sparse positive class, so the gain is not confined to separating the abundant inactive intervals. FunFormer-20k had the highest median AUPRC in all six species and the highest mean AUPRC in five. The exception is chicken, the weakest genome for every model, where FunFormer-1k leads FunFormer-20k on mean AUPRC by a negligible margin (0.538 against 0.535) and on median AUROC by 1.3 pp (0.876 against 0.863). FunFormer compresses long input sequences into a fixed set of tokens, so its context window can be enlarged with little additional cost (Methods).

To choose the window, we swept context length and learning rate in human, a grid of 20 runs evaluating context up to 50 kb (Figure 2c). We selected the 20 kb context on the validation split, and this configuration, FunFormer-20k, is used in all subsequent analyses. The test AUROC of the same grid gives the same picture, 0.968 at 20 kb with no gain at 30 or 50 kb.

### 2.3 The long-context advantage decays with divergence and reverses at the tetrapod-fish split

Within-species accuracy establishes that FunFormer-20k is a strong predictor in every genome, and we next asked how far that accuracy travels. Each within-species model was applied to the genome of every other species and scored on matched (assay, tissue) channels by AUROC (Figure 1b). Which channels those are differs by pair, because the six track sets do not cover the same assays and tissues. Pig, cattle and chicken share a full grid of 40 combinations, human shares 14, and the two fish share 6 to 10 with any partner (Supplementary Fig. S1). The primary analysis uses 23 source-target directions spanning four divergence times, from the pig-cattle split to the tetrapod-fish split. They fall into the three groups defined above, mammal-mammal (6 directions), mammal-chicken (3) and tetrapod-fish (14), the last holding two chicken-fish directions alongside twelve between a mammal and a fish. Directions with chicken as the source are held out, because chicken is the weakest within-species model (Supplementary Table S1) and a poor source confounds the effect of divergence with the quality of the model being transferred. These directions are reported separately in Supplementary Table S2 (Methods). The two salmon-zebrafish directions are also held out, because they form a single pair at 206 Mya that matches none of the three bins (Supplementary Table S2, Methods).

Transfer accuracy declined with divergence time for every model (Supplementary Fig. S2), as it does for human- and mouse-trained models applied to livestock and a single fish species [18]. FunFormer-20k transferred well among mammals, with a mean matched-channel AUROC of 0.888, fell to 0.805 for transfer into chicken, and to 0.650 across the tetrapod-fish split. Referenced to the target species’ own model on the same channels, that is a loss of 5.1 pp among mammals and 5.9 pp into chicken but 28.2 pp across the tetrapod-fish split, or 95%, 93% and 70% of the target model’s own accuracy retained (Supplementary Fig. S3). The low absolute figure for chicken therefore reflects how hard that genome is for any model, its own included. Only the fish drop is a failure of transfer itself. The long-context model and the two 1 kb references, however, decayed at different rates. Among mammals FunFormer-20k led both the best published baseline (0.860) and FunFormer-1k (0.855) by about three points, but it lost accuracy roughly 1.8 times faster with divergence and finished below both at the fish distance (0.650 against 0.728 for both, which are no longer separable there). The two fitted lines cross at 197 and 208 Mya, in the gap between 94 and 320 Mya where none of the 23 primary directions falls. Fitting the same regression on all 359 individual matched channels rather than on the 23 direction means gives a similar slope for the margin (*−*0.028 against *−*0.032 AUROC per 100 million years) with a weaker correlation (*r* = *−*0.61 against *−*0.76), and the same holds for absolute AUROC (Supplementary Fig. S2b). Long context gains the most accuracy where regulatory information is shared and the least where it is not.

The margin of FunFormer-20k over FunFormer-1k isolates this effect, since the two models share architecture and parameter count and differ only in the sequence they see. The margin was +3.4 pp among mammals, +0.4 pp for transfer into chicken, and *−* 7.8 pp across the tetrapod-fish split (Figure 3a, Table 1). The long-context advantage therefore persists only marginally across the amniote split and reverses sign between 320 and 435 Mya. The 23 directions are not independent observations, since they are drawn from six genomes related by a tree and span only four distinct divergence times, and directions sharing a species are correlated. We therefore assessed significance by relabeling the six species onto the tree and recomputing every divergence time and bin, enumerated exactly over all 6! = 720 relabelings (Figure 3b, Methods). The observed trend sits close to the most extreme value the design can produce, since only 12 of the 720 relabelings are at least as extreme and the most extreme reaches *r* = *−* 0.767 against the observed *−* 0.764 (*P* = 0.017, the same for the mammal-versus-fish contrast). The gradient is robust to the choice of 1 kb reference and to the assay set. Measured against the best published baseline, or against the best of all four 1 kb models, the bin margins and species-level *P* are essentially unchanged, and excluding chromatin accessibility makes the gradient slightly steeper rather than shallower (Supplementary Table S3, Supplementary Fig. S4).

**Table 1:**
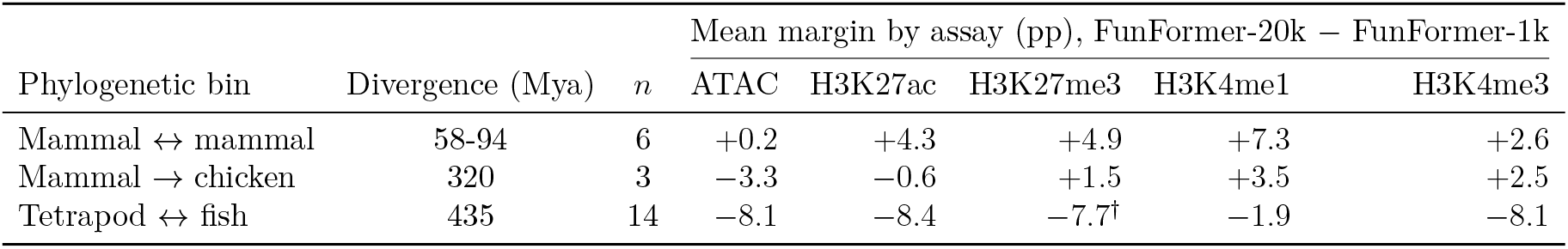
Cross-species transfer margin by phylogenetic bin and assay. Margin is FunFormer-20k minus FunFormer-1k in percentage points, so each entry reflects context length alone; *n* is directions per bin. In the tetrapod-fish bin H3K27me3 and H3K4me1 rest on 12 salmon-only channels, zebrafish lacking those assays; the other three span both fish (35 channels). ^*†*^The H3K27me3 entry rests on two halves that are not comparable, the surviving salmon target tracks differing widely in positive count, and should be read as indicative only.

|  |  |  | Mean margin by assay (pp), FunFormer-20k − FunFormer-1k |  |  |  |  |
|---|---|---|---|---|---|---|---|
| Phylogenetic bin | Divergence (Mya) | *n* | ATAC | H3K27ac | H3K27me3 | H3K4me1 | H3K4me3 |
| Mammal ↔ mammal | 58-94 | 6 | +0.2 | +4.3 | +4.9 | +7.3 | +2.6 |
| Mammal → chicken | 320 | 3 | −3.3 | −0.6 | +1.5 | +3.5 | +2.5 |
| Tetrapod ↔ fish | 435 | 14 | −8.1 | −8.4 | −7.7<sup>†</sup> | −1.9 | −8.1 |

**Figure 3:**
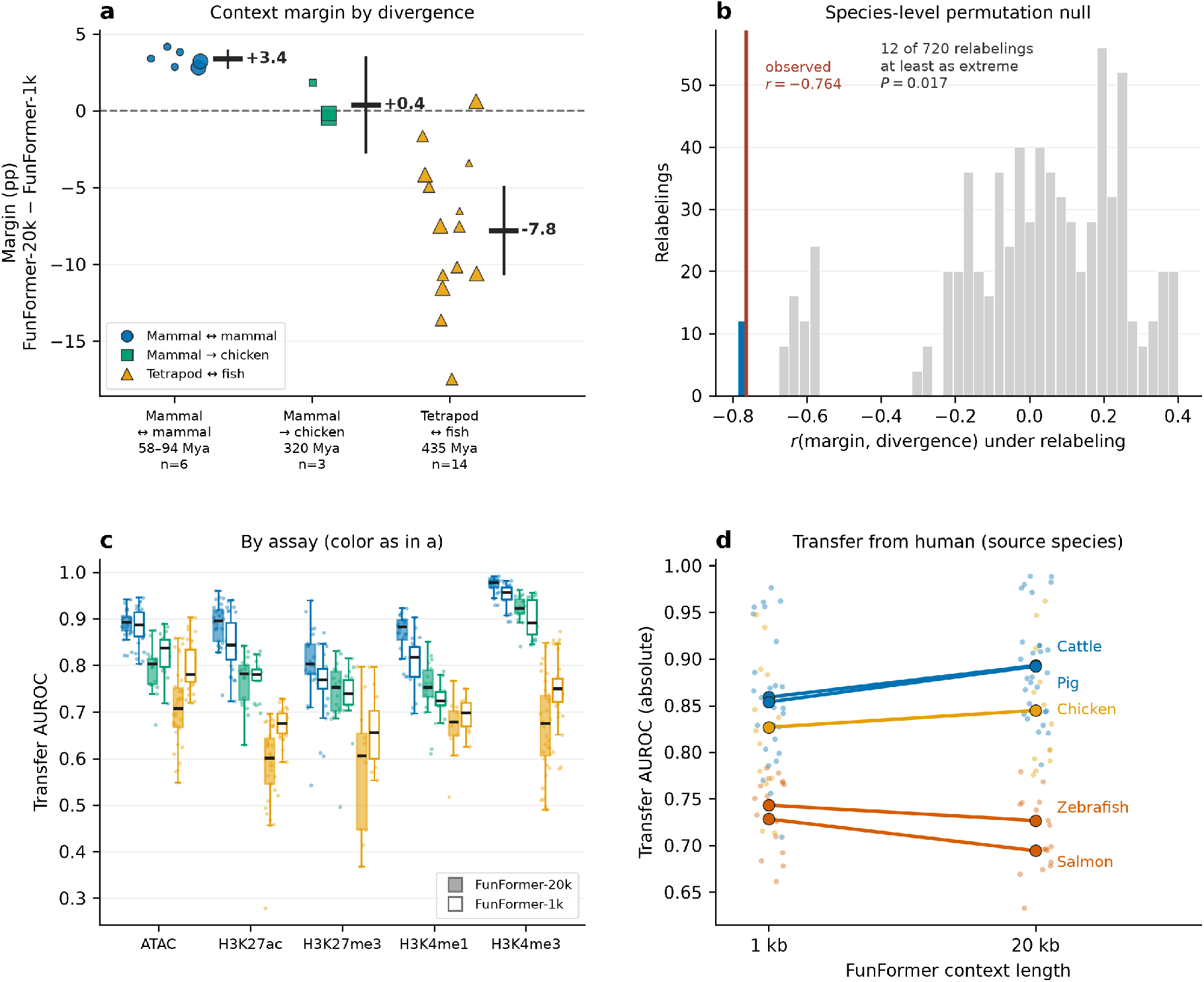
The long-context advantage decays with divergence and reverses at the tetrapod-fish split. **(a)** Context margin, FunFormer-20k minus the parameter-matched FunFormer-1k, by phylogenetic bin. One point per direction, jittered and sized by its 8 to 40 matched channels; bar and whisker, the bin mean and its 95% confidence interval. Bins rather than continuous divergence time, because only four divergence values exist (Supplementary Fig. S2). Within each bin, channels spread more within a direction than between directions (standard deviation 3.74 against 0.54 pp among mammals, 4.34 against 1.24 pp into chicken and 5.64 against 4.92 pp across the tetrapod-fish split). **(b)** All 6! = 720 relabelings of the taxa onto the tree, margins fixed. The observed *r* = *−* 0.764 is marked, the 12 at least as extreme shaded, *P* = 0.017. **(c)** Paired boxes per assay and bin, filled for FunFormer-20k and open for FunFormer-1k, every matched channel a point. Boxes rest on 12 to 35 channels; in the tetrapod-fish bin H3K27me3 and H3K4me1 are salmon-only, zebrafish lacking both. **(d)** Transfer from human as sole source at 1 and 20 kb, one line per target. Absolute AUROC, Supplementary Fig. S2; the margin against the best published baseline, Supplementary Fig. S4.

The reversal is not uniform across regulatory classes, and absolute transfer and long-context margin pick out different assays (Figure 3c, Table 1). By absolute transfer AUROC, the promoter mark H3K4me3 is the most conserved class across amniotes, the same ordering found with human- and mouse-trained models scored against FAANG data [18]. FunFormer-20k transfers it almost unchanged from mammals (0.972) to chicken (0.923) before it collapses to 0.667 at the tetrapod-fish split. Chromatin accessibility is the most uniform class across the whole tree, declining only from 0.888 to 0.792 to 0.710 over the three bins, and it overtakes H3K4me3 at the fish distance. By long-context margin the ranking is different, and the enhancer mark H3K4me1 carried the largest margin among mammals (+7.3 pp) and was the only assay whose margin stayed near zero at the fish distance (*−* 1.9 pp). Every other assay fell to between *−* 7.7 and *−* 8.4 pp there, and accessibility, which carried the smallest margin in both amniote bins, sits mid-range at the fish distance (*−* 8.1 pp). The promoter mark thus transfers best across amniotes, but the extra context helps enhancer-mark prediction for longest.

Restricting to a single source species shows the same gradient without any averaging across directions (Figure 3d). For human as source, moving from 1 kb to 20 kb of context gained 3.8 pp into cattle and 3.4 pp into pig, 1.8 pp into chicken, and lost 1.6 pp into zebrafish and 3.4 pp into salmon. Extending the context amplifies whichever direction the transfer already has, consistent with long context sharpening lineage-specific regulatory grammar rather than capturing a universal syntax. That the two fish behave alike is the key point for interpretation. Both are teleosts carrying the teleost-specific genome duplication, but only salmon carries the further salmonid-specific round, so the reversal is not explained by that duplication. Collectively, these analyses show that the transferability of a regulatory sequence model tracks evolutionary distance, that long context accelerates the loss of transfer with divergence and reverses it beyond the amniote split, and that promoter-associated marks extend the reach of transfer farthest across amniotes.

### 2.4 The long-context advantage is carried by flanking sequence whose target-specific value mostly disappears across the fish boundary

The decline in Figure 3 shows that the value of long context falls with divergence, but not what the model draws from the extra sequence. We addressed this with two analyses on the primary transfer directions, both scored on the same matched channels. The first splits the existing predictions by distance to the nearest transcription start site, and because it reuses predictions already made for the main cross-species analysis we report effect sizes with confidence intervals and no hypothesis tests. The second is an *in silico* flank intervention that replaces the flanking sequence and reruns the model, with its conditions, donor-matching rule and verification criteria fixed before the runs (Methods).

To see where along the genome the long context helps, we split test intervals into those within 2 kb of a protein-coding transcription start site (proximal) and those more than 10 kb away (distal), and computed the context margin separately for each (Figure 4a). Among mammals the margin is positive in both groups and slightly larger proximally (+3.38 pp against +2.66 pp). Across the tetrapod-fish split both are negative, and the proximal loss is nearly three times the distal one (*−* 11.94 pp against *−* 4.12 pp). The absolute values show that this loss comes from FunFormer-20k falling rather than FunFormer-1k improving, at 0.627 against 0.746 proximally and 0.686 against 0.727 distally (Supplementary Table S4). The proximal-distal contrast is nearly the same in both transfer directions, +8.0 pp out of fish and +7.6 pp into fish, so the two are pooled. Proximal and distal values should be compared within a bin and not across bins, because each species holds out a different test chromosome and these differ in gene density, from 88% of human test intervals lying more than 10 kb from a transcription start site to 50% for chicken.

**Figure 4:**
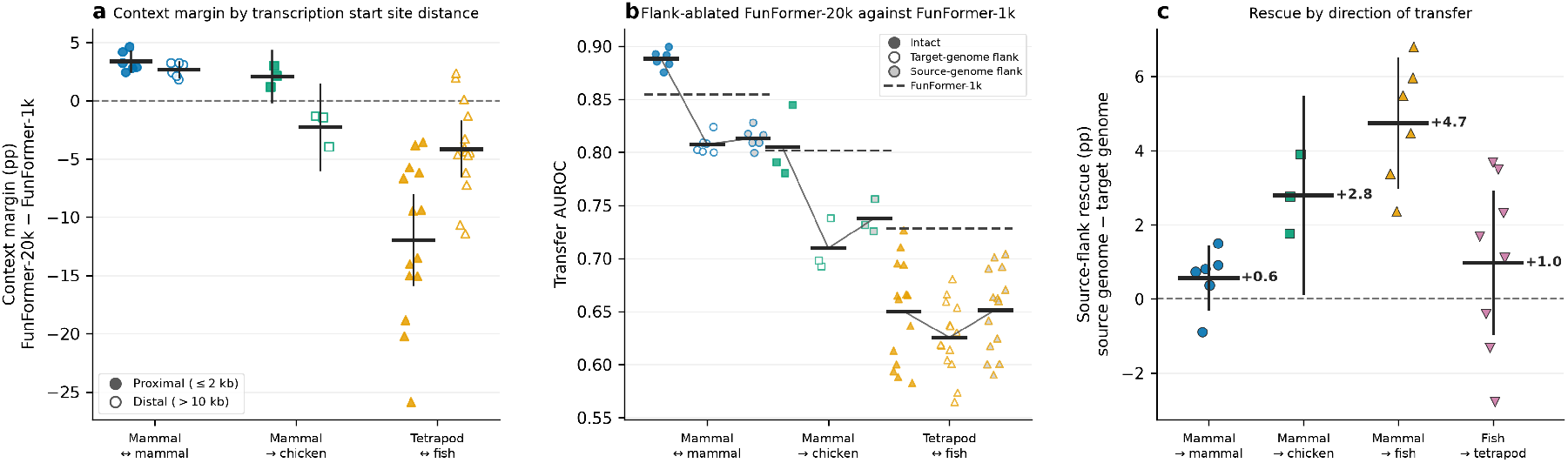
What the long context is used for, and where it stops working. **(a)** Context margin by phylogenetic bin, split by distance from the nearest protein-coding transcription start site. Filled, proximal (*≤* 2 kb); open, distal (*>*10 kb). One point per direction, bar and whisker the bin mean and its 95% confidence interval. **(b)** Matched-channel transfer AUROC when the flanking sequence outside the central 1 kb is replaced, one point per direction. Filled, intact; open, a real GC-matched flank from the target genome; gray, a real GC-matched flank from the source genome. The thin line joins the three means and the dashed line is FunFormer-1k on the same channels, so the question the panel asks is whether an ablated 20 kb model still beats a model that never had a flank. **(c)** Source-flank rescue, AUROC with a source-genome flank minus AUROC with a target-genome flank, by direction of transfer. Panels b and c use the same experiment, and c is the gap between two of the conditions in b rather than an independent measurement.

The flank intervention keeps the central 1 kb of every test interval, replaces the surrounding 19 kb and scores FunFormer-20k under three conditions (Figure 4b). In the intact condition the flank is the locus’s own sequence, whereas in the target-flank condition it is a real sequence of matched length and GC content drawn at random from the target genome, and in the source-flank condition the same kind of sequence is drawn from the source genome instead. The two differences between these conditions answer separate questions, since the value of the flank, intact minus target flank, measures whether the particular sequence around a locus carries information beyond its composition. The source-flank rescue, source flank minus target flank, measures whether the model needs the target’s own sequence at all or only a flank of the kind it was trained on (Figure 4c).

Replacing the flank with a random target-genome sequence costs 8.1 pp among mammals and 9.6 pp into chicken, which shows that the particular flank of a locus carries information. It also leaves the 20 kb model below FunFormer-1k by *−* 4.69 and *−* 9.18 pp respectively, so a long-context model deprived of its context is worse than one built without it. Across the tetrapod-fish split the same replacement costs only 2.4 pp, with 2.8 pp into fish and 2.2 pp out of fish, consistent with little usable flank information being left to remove. A dinucleotide shuffle of the interval’s own flank, which preserves its composition exactly and destroys only the order of the sequence, costs 8.2 and 9.2 pp in the two amniote bins and agrees with the random replacement in sign in every bin and in 18 of 23 directions. In the fish bin the shuffle depends on direction, costing 6.6 pp into fish against 1.7 pp out of it, so its pooled 3.8 pp averages two different halves.

Among mammals, a source-genome flank is worth no more than a target-genome one (+0.58 pp, 95% CI *−* 0.27 to +1.42), so what the model loses is specific to the locus rather than to the genome. When a mammal model is applied to a fish genome, a source-genome flank rescues +4.75 pp (+3.00 to +6.50) and scores 1.97 pp above the intact flank on average (*−* 1.96 to +5.90), so the target’s own flank gives the model no advantage over an arbitrary flank from its training genome. The two human-source directions are the exception, with the intact flank 2.2 and 3.0 pp above the source flank. In the reverse direction, a fish model applied to a tetrapod genome, the rescue is not distinguishable from zero (+0.98 pp, *−* 0.95 to +2.91) and the source flank scores 1.22 pp below intact (*−* 3.65 to +1.21), which leaves the two conditions 0.15 pp apart on average across the whole bin. Rescue depends on which genome supplies the flank rather than on divergence time alone, being larger into chicken (+2.80 pp) than across the tetrapod-fish bin pooled over both directions (+2.59 pp), and within that bin it appears only when the target is a fish. We report the two intervals, rescue present when the target is a fish and absent among mammals, rather than testing the gap between them, because the permutation null used for the main cross-species analysis asks whether a quantity is aligned with the tree, which is not what rescue is claimed to be (Methods).

Taken together, the model reads the flank, draws slightly more from it near transcription start sites among mammals, and, when a mammalian model is applied to a fish genome, on average extracts nothing from the target’s own flank that it cannot get from an arbitrary stretch of the genome it was trained on. Flank value split by the same proximal and distal groups is shown in Supplementary Fig. S5.

### 2.5 Zero-shot variant effect prediction transfers into pig but not into human

We finally asked whether the same phylogenetic pattern carries over to variant-level prediction, a task where non-model species are limited by the scarcity of species-matched training data. We scored eQTL fine-mapping variants with per-track VEP features and trained a random forest to separate causal from non-causal variants (Methods), in human across 49 GTEx tissues [31] and in pig across 19. In pig the long-context model was the best single predictor. It exceeded all three 1 kb baselines trained on the same genome by +2.73 to +3.38 pp, every difference excluding zero (DeepATT +3.38 pp, 95% CI +1.26 to +5.51; DeepSEA +3.20 pp, +0.80 to +5.61; DeepFormer +2.73 pp, +0.24 to +5.22), and it beat each baseline in 16 to 17 of the 19 tissues. The architecture- and parameter-matched control locates most of that gain in the context window, since FunFormer-1k trailed FunFormer-20k by *−*1.98 pp on pig eQTLs (*−*3.46 to *−*0.50) and by *−*2.82 pp on pig sQTLs (*n* = 18 tissues, *−*5.61 to *−*0.03).

In human the long-context model held only a small advantage, leading the three published baselines by +0.28 to +0.65 pp with two of the three intervals excluding zero. It was indistinguishable from its own architecture at 1 kb, where FunFormer-1k scored +0.10 pp above FunFormer-20k (*−* 0.35 to +0.56) and was ahead in 29 of the 49 tissues. The downstream context effect therefore follows the within-species one, since human gains little from context in regulatory-element classification, second least of the six species (+1.00 pp, Supplementary Table S1), and nothing at all in variant prioritization, while pig gains most (+3.56 pp) and retains a clear effect downstream. This is a statement about the present formulation, which scores a per-track change in binary activity at the variant within a 20 kb window and makes no assignment of variants to genes. It is not comparable to long-range regression models, which span about 200 kb to 1 Mb and aggregate predicted effects across the window [4, 9, 11]. Gains from context reported in human for those models concern a task this design does not attempt.

Zero-shot transfer was strongly target-dependent, and this is the informative comparison. Applied to the *data-rich* human target, every foreign model was clearly worse, between *−* 3.61 pp (cattle) and *−* 6.24 pp (zebrafish), with all five intervals excluding zero and winning at most two of the 49 tissues. Applied to the *data-limited* pig target, three of the five foreign models were statistically indistinguishable from the pig-trained model, with zebrafish *−* 0.47 pp (CI *−* 2.81 to +1.88), cattle *−* 0.72 pp (*−* 2.57 to +1.13) and chicken *−* 1.12 pp (*−* 3.76 to +1.52). Human (*−* 1.42 pp, *−* 2.70 to *−* 0.14) and salmon (*−* 1.91 pp, *−* 3.56 to *−* 0.25) were modestly but significantly worse. A model trained on another genome therefore predicts pig eQTLs about as well as one trained on pig. Four of the five foreign 20 kb models also matched or exceeded the best of the three published 1 kb baselines trained on pig itself, taken per tissue, by up to +1.21 pp (zebrafish), and salmon alone fell short, by 0.23 pp. In pig, a foreign long-context model is therefore usually as useful as a native 1 kb one. Zero-shot accuracy declined with divergence time in the human target (Pearson *r* = *−* 0.58 over five source species), in the same direction as the classification gradient, but not in the pig target (*r* = *−* 0.13), where the five sources span only 0.014 AUROC in total. With five sources and so narrow a range, this is a descriptive observation rather than a test. The same two comparisons on pig splicing QTLs (Figure 5c), an independent phenotype over a separate variant set of 18 tissues and 45,897 matched pairs, reproduce both results. FunFormer-20k exceeded all three 1 kb baselines by +2.80 to +3.23 pp, again with every interval excluding zero, and four foreign models (zebrafish, cattle, chicken and human) were indistinguishable from the pig-trained one, salmon alone being worse. On eQTLs human is also worse, so the agreement between the two phenotypes is in the ordering rather than in every interval.

**Figure 5:**
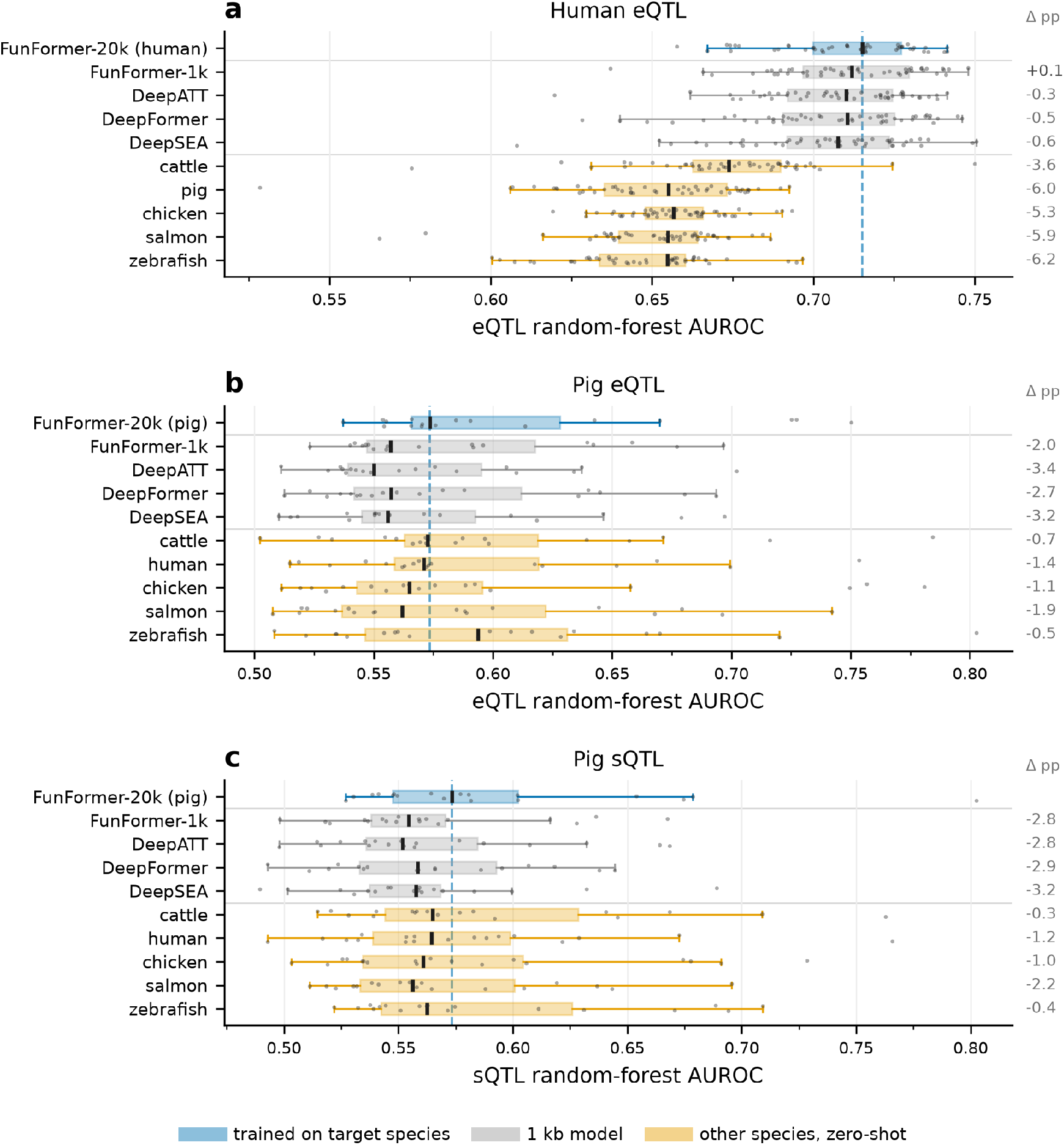
Variant-effect prediction, shown as the distribution across tissues. Random-forest AUROC per feature set, box and whiskers as in Figure 2, every tissue a point. **(a)** Human eQTLs, 49 GTEx tissues. **(b)** Pig eQTLs, 19 tissues. **(c)** Pig splicing QTLs, 18 tissues, an independent phenotype and variant set. Rows are grouped by model access, from training on the target species, through 1 kb models on it (three published baselines and FunFormer-1k), to another species applied zero-shot. Compare rows by theΔpp column, the mean paired difference from the species-matched model, whose median is the dashed line. Boxes show the unpaired distribution across tissues, so a row can sit right of the line while losing in most tissues. Zebrafish in **(b)**, for example, is *−*0.5 pp paired yet ahead in only 6 of 19.

## 3 Discussion

Local and long-range regulatory information, as these models learn it, turned out to have very different phylogenetic reach. A model restricted to the central 1 kb still transferred across 435 million years, reaching a mean matched-channel AUROC of 0.728 across the tetrapod-fish split. The long-range information added by the remaining 19 kb was worth 3.4 pp among mammals separated by up to about 100 million years, was close to zero by the amniote split at 320 Mya, and cost 7.8 pp beyond it. Regulatory element divergence is well documented within mammals [5, 32] and between duplicated salmonid genomes [6], and what the comparison of matched 1 kb and 20 kb models adds is that the long-range information these models learn around a locus has a phylogenetic reach of its own, shorter than that of the local sequence. A long-context model paired with its short-context control is therefore in effect an assay for long-range regulatory information, and the distance at which the long-context advantage disappears estimates where that information stops being shared.

Enlarging the context window amplified whatever direction the transfer already had. Going from 1 to 20 kb with architecture and parameter count unchanged widened both the gains among close mammalian relatives and the losses at the fish distance, and did so for both fish alike. If long context captured a regulatory syntax shared across vertebrates, more of it should help transfer regardless of distance, so the pattern indicates instead that the long-range information the model learns is largely specific to the lineage it was trained in. A model that reads more surrounding sequence has more opportunity to fit features of the training genome that evolve quickly, and it pays for that fit in genomes where those features have changed.

The *in silico* flank intervention tested that account on predictions that did not exist when it was formed and under conditions fixed before the runs. The flank is read, because replacing everything outside the central 1 kb with real, GC-matched sequence cost around 8 pp among mammals and left the 20 kb model below its own 1 kb control, and the informative contrast is where the substitute flank comes from.

Among mammals a flank from the source genome was worth no more than one from the target, so what the model loses is specific to the locus rather than to the genome. When a mammalian model was applied to a fish genome, a source-genome flank on average performed as well as the intact flank, so at that distance the target’s own flank gave the model no more than an arbitrary flank from its training genome. The effect is directional, measurable when transferring into fish rather than out of it, and is estimated as two separate intervals rather than as a test of the gap between them.

Data quality is the obvious alternative to evolutionary distance here, since the more distant species in this study also tend to have smaller track sets and less mature reference assemblies, and zebrafish separates the two. Within-species accuracy is a direct measure of how learnable a species’ own dataset is, and it is sensitive enough to detect a broken one, since built against the wrong assembly every zebrafish model sat near chance (Methods). Built correctly, zebrafish has the third-highest within-species AUROC of the six species, 0.950, above cattle and pig. Transfer into that same genome from a mammal source nonetheless reaches only 0.61 to 0.73, and 20 kb of context is worth *−* 5.6 pp there against the matched 1 kb model. A genome whose own model predicts it at 0.950 cannot be dismissed as too noisy to be predicted, so what fails is transfer from another lineage, not the target data. The two failure modes separate in the other direction as well. Chicken, whose dataset is the weakest of the six (0.876), fails as a *source*, since every chicken-source direction is negative against both 1 kb references (Supplementary Table S2), which is why chicken is used target-side only. As a *target* at the same phylogenetic distance chicken is unremarkable, the three mammal-to-chicken directions spanning *−* 0.5 to +1.8 pp. Weak data therefore degrades the model a species can train, while evolutionary distance degrades the model another species can apply, and zebrafish shows the second without the first.

Within species, the long-context model outperformed every 1 kb model in all six vertebrates, including the three published binary classifiers by 1.0 to 3.8 pp of mean AUROC, and the architecture-matched control places that gain in context rather than architecture, since at 1 kb FunFormer is no better than existing models. Fun-Former is therefore useful in its own right as a within-species predictor, and its fixed token count is what makes 20 kb of context practical for a binary classifier. Across species, the results agree with what has recently been reported for models carried out-side Mammalia. Chromatin annotations transfer at comparable accuracy into cattle, pig and chicken and far less well into the one fish tested, with H3K4me3 transferring best [18], and we recover both orderings on overlapping FAANG data, with each target scored against a model trained on its own genome rather than in absolute terms alone. The claim here is not that transfer declines, which that study and work in plants [19] already establish, but that the value of long-range information declines faster than transfer itself and changes sign. That distinction also resolves an apparent discrepancy, since Enformer, with about 200 kb of input sequence, transfers better than the 1 kb DeepSEA in that study. The two models differ in architecture, output type and training scale, and are trained on human and mouse jointly rather than on one genome, and joint training alone improves held-out accuracy in both species [12], so that comparison cannot attribute the difference to context. FunFormer-20k and FunFormer-1k differ in pooling alone, at identical parameter count and on identical training data, which is what allows the effect measured here to be assigned to context length. A separate line of work concludes that regulatory grammar is more conserved than the sequence carrying it [21]. That is not in conflict either, because that conclusion concerns local regulatory grammar, which in our data also transfers into fish, whereas the boundary measured here concerns the long-range information around it.

The boundary was also not uniform across assays or along the genome, and the absolute and relative views of that heterogeneity answer different questions. By absolute transfer accuracy, the promoter-associated mark H3K4me3 was the most conserved, as it is for human- and mouse-trained models [18], while chromatin accessibility was the most uniform assay across the full range tested. By margin, it was the enhancer-associated mark H3K4me1 whose advantage from long context persisted furthest. Along the genome, intervals within 2 kb of a transcription start site were where long context was the greatest liability across the tetrapod-fish split, costing *−* 11.94 pp against *−* 4.12 pp at distal intervals. Local information near transcription start sites thus transfers well while the long-range information the model reads around them does not. Which sequence features carry that long-range information is a question this design cannot answer, and motif-level attribution could.

For model reuse the boundary gives a practical rule, since in species without their own epigenomic data a model from a close relative benefits from long context, whereas across a deep split a short-context model is the safer choice. In variant scoring, a model trained on one genome scored eQTLs in a data-limited target genome about as well as a model trained on that target, while in a data-rich target every foreign model was clearly worse. This asymmetry does not follow the phylogenetic boundary, since zebrafish, beyond it, transferred into pig as well as any source. It is consistent instead with how well the target is already served by its own model, a well-powered target having little to gain from borrowing any source, close or distant, although two targets cannot separate data availability from other differences between the human and pig benchmarks. A model trained on another genome is therefore a usable variant-effect predictor for a data-limited species. Two features of the analysis set its scope, the first being that it scores a per-track change in binary regulatory activity at the variant itself, over a 20 kb window and with no assignment of variants to genes. It therefore does not measure what long-range regression models measure when they aggregate a predicted effect at a target gene across hundreds of kilobases. The second is that the saving applies to the sequence model only, since fine-mapped variants in the target species are still required to train and evaluate the downstream classifier.

Long-context sequence-to-function models developed for the regression paradigm, most of them trained on human data with a smaller mouse component [4, 9, 11], are increasingly proposed as general-purpose tools for interpreting the noncoding genome across species. The boundary identified here was measured for a binary classifier, and we hypothesize that the same amplification by context length operates in regression models trained on similarly deep context, which would bound their cross-species reach as well. Testing that directly, most naturally in binary terms on matched regulatory channels as done here, is an open question raised rather than answered by this work.

The design bounds how precisely the boundary can be placed. The six species span four distinct divergence times, three of the twenty-three primary directions fall in the mammal-chicken bin, and that bin rests on transfer into chicken rather than out of it, so the transition is located between 320 and 435 Mya rather than resolved within that interval. Both fish carry the teleost genome duplication, which coincides with the tetrapod-fish split, so this design cannot separate it from divergence time. Species inside the gap would narrow it, among them amphibians at about 350 Mya from mammals and the lobe-finned fishes such as coelacanth or lungfish at about 415 Mya, and a second bird would supply the avian analogue of the mammal-mammal bin and establish whether a bird source transfers at short range at all. Our margins also describe the typical locus on a held-out chromosome. Regulatory blocks around developmental genes keep their synteny from teleosts to mammals [7], and duplicated salmonid enhancers carrying deeply conserved noncoding elements are preferentially retained [6], so such blocks may keep long-range information beyond the boundary, but our test chromosomes contain too few of them to resolve this. Motif-level or attention-based attribution of what the model reads at 20 kb but not at 1 kb would identify the sequence features behind the long-range information and test whether they are the rapidly evolving features proposed here. Whether light fine-tuning on a small amount of target-species data can rescue transfer beyond the boundary, which initializing a zebrafish model from mammalian weights suggests is plausible [22], and whether joint multi-species training encourages a more universal regulatory grammar than single-species training does, are both directly testable extensions of the pipeline built for this study.

Overall, comparing matched long- and short-context models across six vertebrates shows that the long-range information these models learn has a shorter phylogenetic reach than local regulatory information. Its value is positive among close relatives, near zero at the amniote split and negative beyond it, so around a typical locus this information stops being shared between vertebrate genomes somewhere between 320 and 435 million years of divergence, well before local sequence stops being informative. Within one genome FunFormer is the strongest regulatory-element predictor among the models tested, but that advantage comes from context and does not carry over indiscriminately across species. Context length in genomic sequence models should therefore be chosen with the intended scope of use in mind, long within one genome or among close relatives and short for reuse across deep splits. For variant-effect prediction, the data-limited target tested could borrow a model from most sources, while in the data-rich target every foreign model lost accuracy.

## 4 Methods

### 4.1 Data

We assembled binary regulatory-track datasets for six vertebrates. Human tracks (423) are from the EN-TEx resource [28], mapped to GRCh38. Pig (80 tracks, Sscrofa11.1), cattle (95, ARS-UCD1.2), salmon (301, Ssal_v3.1) and chicken (97, GRCg6a) are the FAANG- and AQUA-FAANG-derived datasets described in our earlier work [29], drawn from the same consortium resources as the livestock and fish annotations of earlier cross-species work [18]. Zebrafish (37 tracks) is the cis-regulatory element map of Yang et al. [30] (GEO accession GSE134055), mapped to GRCz10, the assembly on which that resource’s peak calls were built; pairing those coordinates with the later GRCz11 assembly decouples the labels from the sequence and drives every model to chance.

Every dataset contains binary peak calls for chromatin accessibility (ATAC-seq; DNase-seq in chicken) and histone marks. H3K27ac, H3K27me3, H3K4me1 and H3K4me3 are present in all species except zebrafish, which has H3K27ac and H3K4me3 only. CTCF and H3K36me3 occur in some species, are not shared across all six, and are excluded from the cross-species comparisons. Track names encode assay and tissue per species and were parsed into (assay, tissue, track index) records; human EN-TEx tracks are identified by ENCODE accession and joined against the EN-TEx sample metadata, with free-text UBERON biosample terms mapped to five canonical tissue labels (Adipose, Liver, Lung, Muscle, Spleen) used consistently across all six species.

Each dataset is stored as a HuggingFace Arrow dataset of predefined genomic intervals with per-track binary labels and train, validation and test splits assigned upstream of this study, prior to any analysis reported here. The input window is centered on each interval’s midpoint and extended symmetrically to the model’s context length; both the forward strand and its reverse complement are retrieved from the genome FASTA at runtime. Positions beyond a chromosome boundary are N-padded rather than dropped, so every window has exactly the requested length.

### 4.2 Model architectures

We compared FunFormer against three 1 kb baselines, all trained with unweighted per-track binary cross-entropy loss on raw logits. The baseline implementations were adapted from the code released with DeepFormer (https://github.com/YZ20211221/DeepFormer). DeepSEA is a three-layer CNN (channels 4 *→*320 *→*480 *→*960, kernel size 8, max-pooling by 4 after the first two layers) followed by a two-layer fully connected classifier [8]. DeepFormer extends this with two convolutional layers per stage and batch normalization, and replaces the final classifier’s input with the output of a flow-attention module over the pooled sequence representation [26]. DeepATT uses a single wide convolution (kernel size 30, 4 *→*1024 channels, max-pooling by 15), a bidirectional LSTM (hidden size 512 per direction) and a four-head cross-attention layer in which each track’s one-hot identity vector is the query, so every track’s logit is read from its own attention-weighted mixture of the sequence [27].

FunFormer uses a three-layer convolutional encoder (channels 4 *→*320 *→*480 *→*512, kernel sizes 15, 9, 9) whose pooling factors are chosen so that any input from 1 to 50 kb is compressed to exactly 500 tokens (a 20 kb input uses pool sizes 5, 4, 2). Those tokens are processed by transformer layers with pre-RMSNorm, query/key RMSNorm, rotary positional embeddings [33], bidirectional scaled-dot-product attention (dispatching to FlashAttention-2 [34] on CUDA) and a SwiGLU feed-forward block [35]. All FunFormer models reported here use a single transformer layer, hidden size 512, 8 attention heads and an intermediate size of 1024. The pooling layers carry no parameters, and the classifier head always receives the same 500 tokens. The head projects each token to 256 dimensions, flattens the tokens and maps them to the tracks through one hidden layer, so every context length has an identical parameter count for a given track set.

### 4.3 Training

All models were trained from scratch within each species using the Hugging Face Trainer, with AdamW, weight decay 0.01, a linear learning-rate schedule with 1% warmup and bf16 mixed precision. Per-device batch size and gradient accumulation were scaled with context length so that the *effective* batch size is 1000 for every model reported here, so no comparison in this study is confounded by batch size. Training used early stopping on validation binary cross-entropy loss (patience 5), with a maximum of 30 epochs for FunFormer and 50 for the 1 kb baselines; early stopping rather than the epoch ceiling determined the stopping point in every FunFormer-1k run, so the difference in ceiling does not confound the comparison. Each training example provides both strands of an interval, one of which is chosen at random each training step, while both are scored and averaged at evaluation. To choose the context length we swept five lengths (5, 10, 20, 30, 50 kb) by four learning rates in human only, a grid of 20 runs, and selected 20 kb on the validation split, and this length, FunFormer-20k, was then used unchanged in every other species. Within each species, every model, the three baselines and both FunFormer configurations alike, was trained at four learning rates, and the run with the best validation performance gave the checkpoint used in all analyses, while the test split was used only to report final performance.

### 4.4 Evaluation

Splits are by whole chromosome and were assigned upstream of this study. Each species holds out one chromosome for validation and one for test and trains on every remaining autosome, so no test interval shares a chromosome with any interval the model was fitted on. The held-out test chromosome is 21 in human, cattle, chicken, salmon and zebrafish and 16 in pig, while validation uses 22 in human, 17 in pig and 25 in cattle, chicken, salmon and zebrafish. The test splits hold 233,549 intervals in human, 399,721 in pig, 349,314 in cattle, 34,224 in chicken, 299,099 in salmon and 229,478 in zebrafish, before the two filters described below.

For each species and model we computed per-track AUROC and AUPRC on the held-out test split; tracks with only one label class in the test set were excluded from that model’s evaluation. The summary values reported in Supplementary Table S1 and Figure 2 are the mean over all evaluated tracks in a species, for the learning rate selected on the validation split for each model. The test split was used only to report final performance.

Two filters are applied identically to every species and to both the within-species and the cross-species evaluation. First, test intervals whose 20 kb window is more than 50% N on either side of the central 1 kb are excluded, since such an interval gives the model a window that is largely unplaced sequence. Second, a track is evaluated only if it carries at least 50 positives among the intervals that survive the first filter; counting positives on the full set would admit a track whose positives all lie in the excluded regions. This leaves 401 of 423 human tracks and 211 of 301 salmon tracks, and the other four species lose none. In the cross-species evaluation the floor is applied to the target track of each replicate pair, which removes 48 tracks across 10 channels but drops no channel entirely.

Neither filter changes the decline of the margin with divergence materially, since the all-assay bin margins move by at most 0.7 pp, the pooled correlation from *r* = *−* 0.758 to *−* 0.764, and the species-level exact *P* not at all. The same twelve relabelings remain at least as extreme as the observed one with and without the filters. Supplementary Table S3 reports the main results without either filter.

For every ordered species pair we applied the source-trained model to the target species’ genome and test intervals, then compared source and target tracks on matched (assay, tissue) channels. Matched channels were identified with every species’ track records restricted to the five shared assays (ATAC, H3K27ac, H3K27me3, H3K4me1, H3K4me3) and assay-tissue combinations intersected pairwise across all fifteen species pairs, after three species-specific restrictions. Chicken DNase-seq tracks are relabeled “ATAC” so that they match ATAC-seq tracks elsewhere, a technology confound that affects only chicken, since every other species here profiled accessibility with ATAC-seq; chicken itself contributes two ATAC-seq tracks alongside fifteen DNase-seq ones. Salmon is restricted to liver and muscle and zebrafish’s two embryonic tissues are dropped, removing tissues no other species profiles. Chicken’s three brain tissues are excluded whenever chicken is matched against human or salmon, neither of which has brain tracks, and retained otherwise. The matched vocabulary therefore differs by pair, from eight tissues among pig, cattle and chicken down to two for any pair involving salmon; Supplementary Fig. S1 gives the full coverage.

Because a species can have several replicate tracks for one (assay, tissue) combination, each matched group stores a list of track indices per species. We computed AUROC for every pairing of a source replicate with a target replicate and averaged those to one number per channel per direction. Three margins are computed for every direction, the first being FunFormer-20k minus FunFormer-1k, which holds architecture and parameter count fixed and is the primary one reported throughout. The second is FunFormer-20k minus the best of the three 1 kb baselines, and the third is FunFormer-20k minus the best of all four 1 kb models, which is the strictest. All-assay margins, which include accessibility, are reported as the main result, and margins excluding accessibility are reported alongside as a robustness check (Supplementary Table S3).

Of the 30 directions computed, 23 form the primary set. The five chicken-source directions are held out because chicken’s within-species FunFormer AUROC (0.876) was the weakest of the six species, a training-quality confound rather than a phy-logenetic signal. Chicken is therefore used target-side only, and those directions are reported as a sensitivity check in Supplementary Table S2. The two salmon-zebrafish directions are held out because they form a single pair whose divergence (206 Mya) falls between the mammal-mammal bin and the amniote split rather than at the tetrapod-fish distance, and they rest on six matched channels. Divergence times were taken from TimeTree [36], with the amniote split matching fossil calibrations [37], giving pig-cattle 58 Mya (range 51 to 65), human-pig and human-cattle 94 Mya (90 to 96), mammal-chicken 320 Mya (312 to 330, the amniote split), and tetrapod-to-either-fish 435 Mya (430 to 450, the Sarcopterygii/Actinopterygii split). Salmon and zebrafish therefore sit at the same divergence from every tetrapod here and differ in the salmonid-specific whole-genome duplication that only salmon carries; they diverged from each other 206 Mya (200 to 230).

### 4.5 Distance to transcription start sites

Transcription start sites are the 5^*′*^ ends of protein-coding transcripts in the Ensembl GTF for the assembly each model was trained on, which for zebrafish means release 91, the last carrying GRCz10. Each test interval is assigned the distance from its midpoint to the nearest such site, strand-agnostic. Proximal is *≤* 2 kb and distal *>*10 kb, with the intervening band dropped. A sensitivity analysis at *≤* 1 kb and *>*20 kb leaves every bin margin the same in sign and ordering. Every stratified quantity is computed on the same N-excluded interval set as the unstratified margins, so the two are directly comparable.

Stratification is applied to positives only, with the negative set shared between strata, so the two AUROCs differ in which positives must outrank the same background rather than in the background itself. A channel enters the analysis only if it clears the 50-positive floor in both strata, so proximal and distal rest on one channel set and are paired; 354 of 359 channels survive. Comparisons are made within a direction, as distal minus proximal.

### 4.6 In silico flank intervention

The flank intervention is a region-level perturbation analysis, in the family of occlusion and global importance analysis [38], in which a defined part of the input is replaced and the change in model output is measured across the whole test set. For each primary direction the central 1 kb of every test interval is kept and the surrounding 19 kb replaced, with FunFormer-20k and the same checkpoints used elsewhere, under one of three replacements. *Shuffled* applies an Altschul–Erickson dinucleotide shuffle [39] to each flank, preserving its length, GC content and dinucleotide composition exactly and destroying only the arrangement. *Random, same genome* substitutes a real sequence drawn from the target genome, barring its test chromosome; for a within-species direction the donor comes from the validation chromosome instead, so the donor is unseen by the model in every case. *Random, source genome* substitutes a real sequence from the source species’ held-out chromosome, again sequence the model was not fitted on.

Donors are matched to the GC content of the flank they replace within 2%, widening to 3% and otherwise keeping the closest candidate found, and are rejected if more than 1% N or containing an N run of 50 bp or more. Achieved GC differences and the donor coordinates are recorded per flank, so a run is reproducible without rerunning the sampler, and N is never used as filler. The reverse strand is recomputed from the modified forward sequence rather than reused, and the intact condition reproduces the predictions of the main evaluation exactly.

### 4.7 Statistics

The decrease of the transfer margin with divergence time was tested three ways across the 23 primary directions, by the Pearson correlation between margin and divergence time, by a Jonckheere-Terpstra ordered-trend statistic across the three phylogenetic bins, and by a one-sided test of the mean margin difference between the mammal-mammal and tetrapod-fish bins. All *P* values are permutation-based rather than asymptotic, given the small sample. The null permutes the six species onto the tree and recomputes each direction’s divergence time and bin from the relabeled assignment, holding the observed margins fixed. It is the directional analogue of a Mantel test and respects the correlation between directions that share a species. With six species there are exactly 6! = 720 relabelings, so the null is enumerated rather than sampled and the *P* value is exact. For the primary margin, the Pearson correlation and the Jonckheere-Terpstra statistic (*J* = 2.0) both give *P* = 12/720 = 0.017. Every test was run for all three margin definitions, with and without accessibility (Supplementary Table S3), and was repeated without the salmon *→*chicken direction as a leave-one-out check, which left the species-level *P* unchanged.

Tests use direction means rather than the 359 raw matched channels, because channels within a direction share a source model, a target genome and a checkpoint. This is the conservative choice. Refitting the margin on the raw channels keeps its sign and most of its slope (*−*0.028 against *−*0.032 AUROC per 100 million years, *r* = *−*0.61 against *−* 0.76), and Supplementary Fig. S2b shows the same for absolute AUROC. For source-flank rescue the three replacement conditions, the donor-matching rule and the verification criteria were fixed before the ablation runs. Donors are drawn once per interval under a single seed, so a direction’s value averages thousands of independent draws, and bin means carry Student *t* intervals over directions. The species-level permutation is not applied to rescue, which is a within-direction contrast between two conditions scored on the same intervals with the same checkpoint rather than a quantity claimed to align with the tree. The splits of rescue by source clade and by distance to a transcription start site are descriptive, and the stratification by transcription start site reuses predictions already made for the main cross-species analysis and is reported without tests.

Every confidence interval in this paper is a two-sided 95% Student *t* interval on a mean,

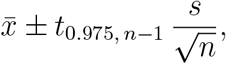

with *s* the sample standard deviation on *n −* 1 degrees of freedom. The *t* quantile is used at every *n* rather than the normal 1.96, which matters most where the sample is smallest. What *n* counts differs between analyses and is the unit of replication in each. For the bin means of Figure 3a and Figure 4a,b it is the transfer direction, giving *n* = 6, 3 and 14 for the three bins, and Figure 4c, which splits the tetrapod-fish bin by direction, has *n* = 6, 3, 6 and 8. For the variant-effect comparisons it is the tissue, *n* = 49 in human and 19 in pig, and the interval is taken on the paired within-tissue difference between two feature sets rather than on either mean separately. For Supplementary Fig. S5 it is the matched channel.

### 4.8 Variant effect prediction and eQTL random forest

For a given trained model, the per-variant, per-track impact score is

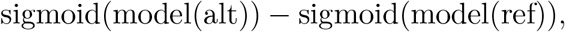

computed for the reference and alternate allele centered in a window of the model’s own training context length (1 kb for the three baselines and FunFormer-1k, 20 kb for FunFormer-20k), averaged over both strands; only single-nucleotide variants are scored. Human variants and their fine-mapped causal and non-causal labels across 49 GTEx tissues [31] are the benchmark VCFs released with the Enformer study [4], one matched pair of files per tissue at a 1:1 ratio. Scoring is restricted to the autosomes, which is what the training intervals cover in every species; this removes 2,240 of the 97,922 human records (2.3%, all on chrX).

Pig variants and labels were built from PigGTEx [40] SuSiE [41] fine-mapping output under the same rule as the Enformer benchmark. Causal variants are those with a posterior inclusion probability above 0.9; each is matched to one non-causal control drawn first from the same gene (posterior below 0.01, |*z*| *>* 4) and, once that pool is exhausted, from any gene tested in the same tissue (posterior below 0.01, |*z*| *>* 6). Controls are sampled without replacement and tissues with fewer than 200 causal variants are dropped, giving 19 tissues and 32,375 matched pairs for eQTLs and 18 tissues and 45,897 pairs for splicing QTLs, written as VCFs in the same format as the human benchmark and scored by the same code. For each tissue we trained a random-forest classifier (100 trees, maximum depth 10, minimum samples per split 10) to separate causal from non-causal variants using the per-track score vector as features, evaluated with ten-seed, 10-fold cross-validation (100 runs) and summarized by the mean AUROC. Feature sets are compared within tissue, by the mean paired difference in AUROC across the tissues they share, with a 95% confidence interval on that paired difference (Statistics).

## Data and code availability

This study analyzed existing, publicly available data. Human tracks are from the EN-TEx resource [28], and cattle, pig, chicken and salmon tracks are from the FAANG and AQUA-FAANG projects, processed as described in our earlier work [29]. Zebrafish tracks are from Yang et al. [30] and are deposited in the Gene Expression Omnibus under accession GSE134055. Fine-mapped eQTL variants are from GTEx [31] for human and from PigGTEx [40] for pig. Reference genome sequences were downloaded from Ensembl [42] (https://www.ensembl.org) as the GRCh38 primary assembly for human and the top-level assemblies Sscrofa11.1 for pig, ARS-UCD1.2 for cattle, GRCg6a for chicken, Ssal_v3.1 for Atlantic salmon and GRCz10 for zebrafish, the last from release 91, the final Ensembl release to carry that assembly. Transcription start sites were taken from the matching Ensembl gene annotations, release 112 for human and salmon, 110 for cattle, 106 for pig, 105 for chicken and 91 for zebrafish. Divergence-time estimates were obtained from Time-Tree (http://timetree.org) [36] and the fossil calibrations cited in Methods. Code implementing FunFormer, the baseline models and the full analysis pipeline is available at https://github.com/SciFarmers/funformer.

## Acknowledgments

The computations, data analysis and model training were performed on resources provided by Sigma2, the National Infrastructure for High-Performance Computing and Data Storage in Norway, using the Olivia supercomputing cluster under project nn9780k. This work received no specific funding.

## Author contributions

Conceptualization, D.T.N.; methodology, D.T.N.; software, B.L. and D.T.N.; formal analysis, B.L. and D.T.N.; data curation, D.T.N., B.L., H.N., L.G. and S.R.S.; visualization, B.L., D.T.N. and O.F.; resources, T.T.T. and O.A.A.; supervision, D.T.N. and T.T.T.; writing – original draft, D.T.N. and B.L.; writing – review and editing, all authors.

## Declaration of generative AI and AI-assisted technologies in the manuscript preparation process

During the preparation of this work, the authors used Claude Code and Claude (Anthropic), with the Claude Sonnet 4.6, Sonnet 5, Opus 4.8, Opus 5 and Opus 5.5 models, to assist with code implementation and to help edit the text during revision. The authors reviewed and edited the output as needed and take full responsibility for the content of the published article.

## Supplementary Information

**Table S1:** Within-species test AUROC and AUPRC for five models across six species. Tracks are those passing the two filters described in Methods, so fewer than the number trained on. Values are means over those tracks for each model at the learning rate selected on the validation split, the best per species bold for each metric. Reading across a row separates the architecture effect (FunFormer-1k against the three baselines) from the context effect (FunFormer-20k against FunFormer-1k). Text differences use unrounded values.

| Species | Tracks | Metric | DeepSEA | DeepFormer | DeepATT | FunFormer-1k | FunFormer-20k |
|---|---|---|---|---|---|---|---|
| Human | 401 | AUROC | 0.951 | 0.956 | 0.957 | 0.957 | **0.967** |
|  |  | AUPRC | 0.349 | 0.374 | 0.377 | 0.370 | **0.426** |
| Pig | 80 | AUROC | 0.882 | 0.889 | 0.889 | 0.892 | **0.927** |
|  |  | AUPRC | 0.352 | 0.369 | 0.372 | 0.379 | **0.470** |
| Cattle | 95 | AUROC | 0.924 | 0.928 | 0.928 | 0.929 | **0.949** |
|  |  | AUPRC | 0.459 | 0.478 | 0.480 | 0.483 | **0.533** |
| Chicken | 97 | AUROC | 0.847 | 0.865 | 0.856 | 0.868 | **0.876** |
|  |  | AUPRC | 0.500 | 0.535 | 0.522 | **0.538** | 0.535 |
| Salmon | 211 | AUROC | 0.931 | 0.937 | 0.942 | 0.943 | **0.955** |
|  |  | AUPRC | 0.308 | 0.337 | 0.355 | 0.361 | **0.395** |
| Zebrafish | 37 | AUROC | 0.905 | 0.925 | 0.925 | 0.929 | **0.950** |
|  |  | AUPRC | 0.426 | 0.516 | 0.509 | 0.530 | **0.584** |

**Table S2:** Chicken-as-source and salmon-zebrafish cross-species transfer margins, a sensitivity check outside the primary 23-direction set. Chicken is used target-side only (Methods), its within-species FunFormer AUROC (0.876) being the weakest of the six, and all five chicken-source directions are negative against both 1 kb references, consistent with a training-quality confound rather than phylogeny. The ordering across targets is also unstable between the two margin definitions, and under neither does it follow phylogeny. Against the published baselines cattle (*−*10.3 pp) sits between the two fish, worse than salmon (*−*9.8 pp) and better than zebrafish (*−*10.8 pp), while against FunFormer-1k cattle (*−*9.0 pp) is worse than salmon (*−*6.7 pp). These directions are therefore excluded from every trend statistic. The two salmon-zebrafish directions, listed below the rule, are excluded because they form a single pair at 206 Mya that matches none of the three bins.

| Target | Divergence (Mya) | FunFormer-20k | FunFormer-1k | Best baseline AUROC | Best baseline | Margin vs baseline (pp) | Margin vs FunFormer-1k (pp) |
|---|---|---|---|---|---|---|---|
| Human | 320 | 0.776 | 0.833 | 0.822 | DeepSEA | −4.6 | −5.7 |
| Pig | 320 | 0.741 | 0.796 | 0.792 | DeepSEA | −5.1 | −5.4 |
| Cattle | 320 | 0.712 | 0.801 | 0.815 | DeepSEA | −10.3 | −9.0 |
| Salmon | 435 | 0.562 | 0.629 | 0.659 | DeepATT | −9.8 | −6.7 |
| Zebrafish | 435 | 0.637 | 0.728 | 0.745 | DeepATT | −10.8 | −9.1 |
| *Salmon–zebrafish directions, also outside the primary set* |  |  |  |  |  |  |  |
| Salmon → zebrafish | 206 | 0.827 | 0.791 | 0.805 | DeepFormer | +2.3 | +3.6 |
| Zebrafish → salmon | 206 | 0.799 | 0.830 | 0.821 | DeepFormer | −2.2 | −3.2 |

**Table S3:**
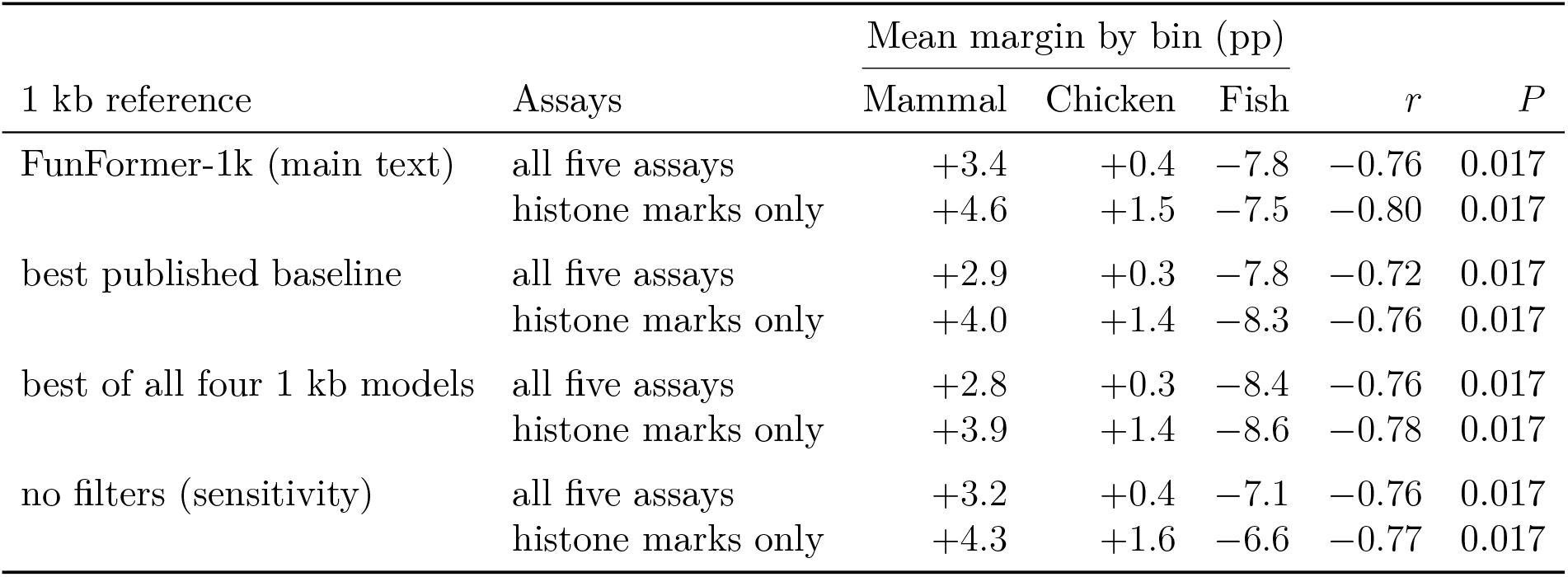
The cross-species gradient is robust to the 1 kb reference and to the assay set. Mean transfer margin by phylogenetic bin for each 1 kb reference, on all five assays and on the four histone marks alone. *r* is the pooled Pearson correlation with divergence time over the 23 primary directions, *P* the species-level exact test over all 6! = 720 relabelings (Methods). Sign pattern, the reversal beyond the amniote split, and significance hold for every reference, assay set and filter setting, and excluding chromatin accessibility steepens the gradient rather than flattening it.

**Table S4:** Absolute AUROC behind the stratified context margin. The margin of Figure 4a is the difference of the two columns per stratum, given here so it can be read against its baseline. *n* is directions per bin. The tetrapod-fish bin is also split by transfer direction. The distal-minus-proximal contrast the figure relies on is +7.6 pp into fish and +8.0 pp out of fish, so pooling the two does not create the effect. At that distance the proximal penalty is FunFormer-20k falling rather than FunFormer-1k excelling.

|  |  | Proximal (≤2 kb) |  | Distal (>10 kb) |  |  |
|---|---|---|---|---|---|---|
| Phylogenetic bin | *n* | FF-20k | FF-1k | FF-20k | FF-1k | Distal − proximal |
| Mammal ↔ mammal | 6 | 0.953 | 0.919 | 0.854 | 0.827 | −0.7 |
| Mammal → chicken | 3 | 0.862 | 0.841 | 0.767 | 0.789 | −4.3 |
| Tetrapod ↔ fish | 14 | 0.627 | 0.746 | 0.686 | 0.727 | +7.8 |
| *into fish* | 6 | 0.639 | 0.739 | 0.687 | 0.711 | +7.6 |
| *out of fish* | 8 | 0.618 | 0.752 | 0.686 | 0.740 | +8.0 |

**Figure S1:**
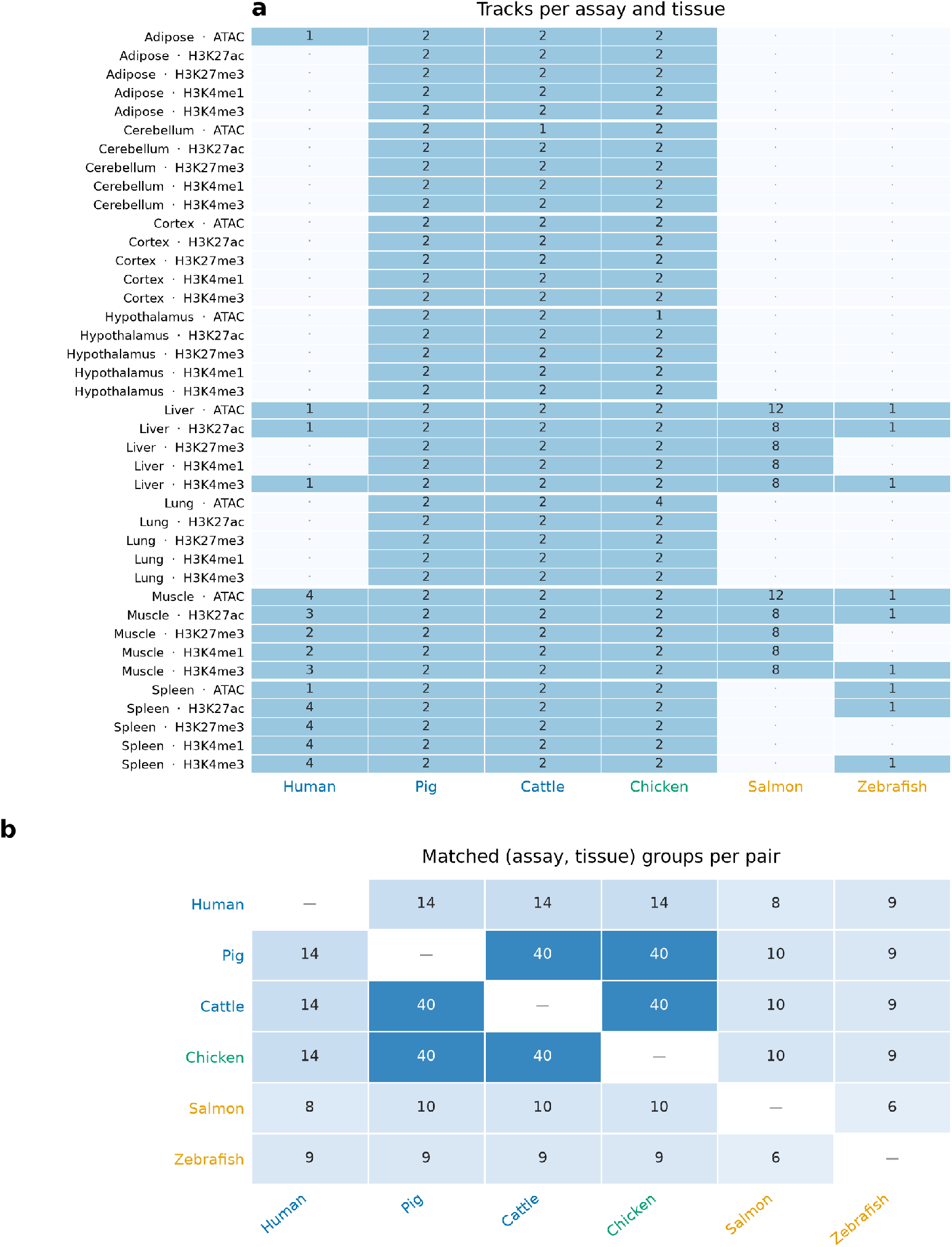
Assays and tissues per species, and matched channels per pair. Transfer is scored only on (assay, tissue) combinations shared by source and target, so the comparison set differs for every pair. **(a)** Tracks per combination and species, for the 40 present in at least two species; a dot marks one the species lacks. Matching is by combination, replicate pairs within one averaged to a single AUROC (Methods). **(b)** Matched groups per pair. Pig, cattle and chicken share the full grid of 40, human 14, the fish 6 to 10 with any partner. Labels colored by clade.

**Figure S2:**
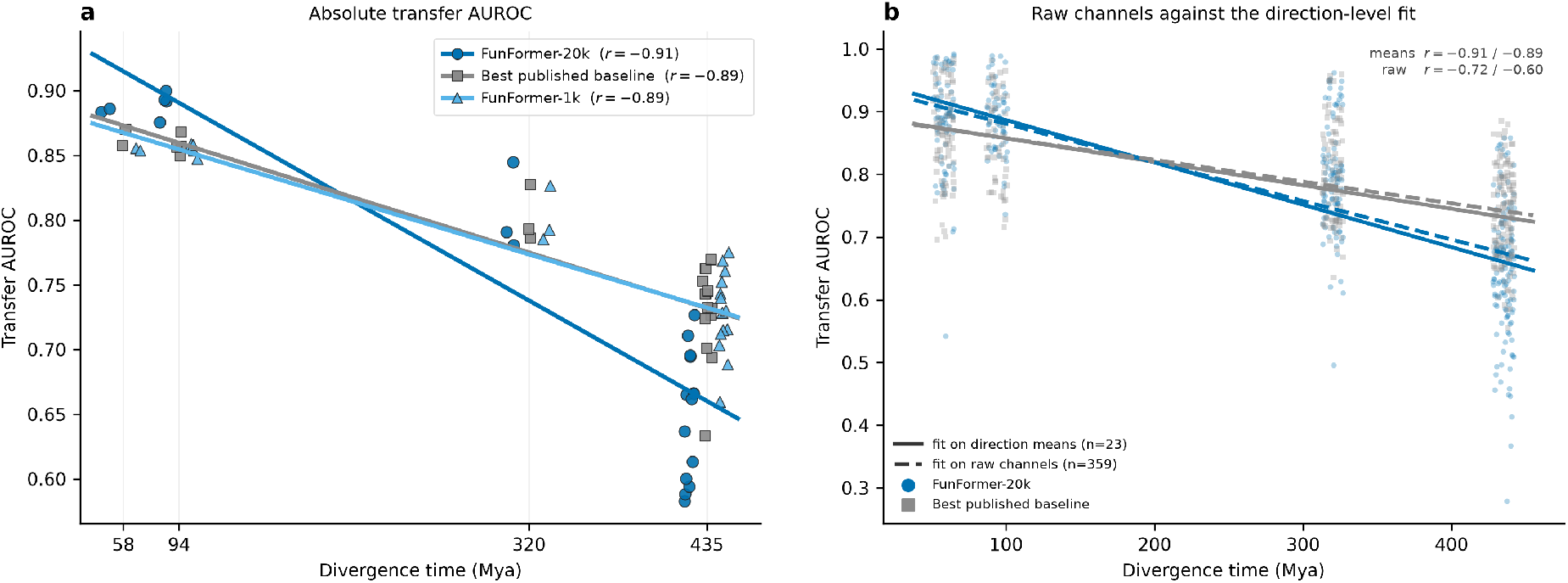
Absolute transfer AUROC, and whether its regression depends on fitting direction means. **(a)** Transfer AUROC against divergence time, one point per direction (*n* = 23), for FunFormer-20k, the best published baseline and FunFormer-1k. The series sit in jittered lanes either side of each divergence value; the fits use the true values. The lines cross at 197 and 208 Mya, inside the unmeasured 94–320 Mya gap, which is why the main figure reports the margin by bin. Absolute AUROC also conflates transfer loss with target-genome predictability (Supplementary Fig. S3). **(b)** The same regression refitted on all 359 raw channels (dashed) against the direction-level fit (solid). Both agree on sign and magnitude, the raw *r* weaker, as expected when channel noise is averaged away. Channels within a direction share a source genome, target genome and checkpoint, so they are not independent and the main analysis averages them first. For the context margin, permuting the 23 direction values directly, without that structure, gives *P* = 0.00005 for its correlation with divergence time (200,000 permutations), against the species-level *P* = 0.017 the main analysis reports (Methods).

**Figure S3:**
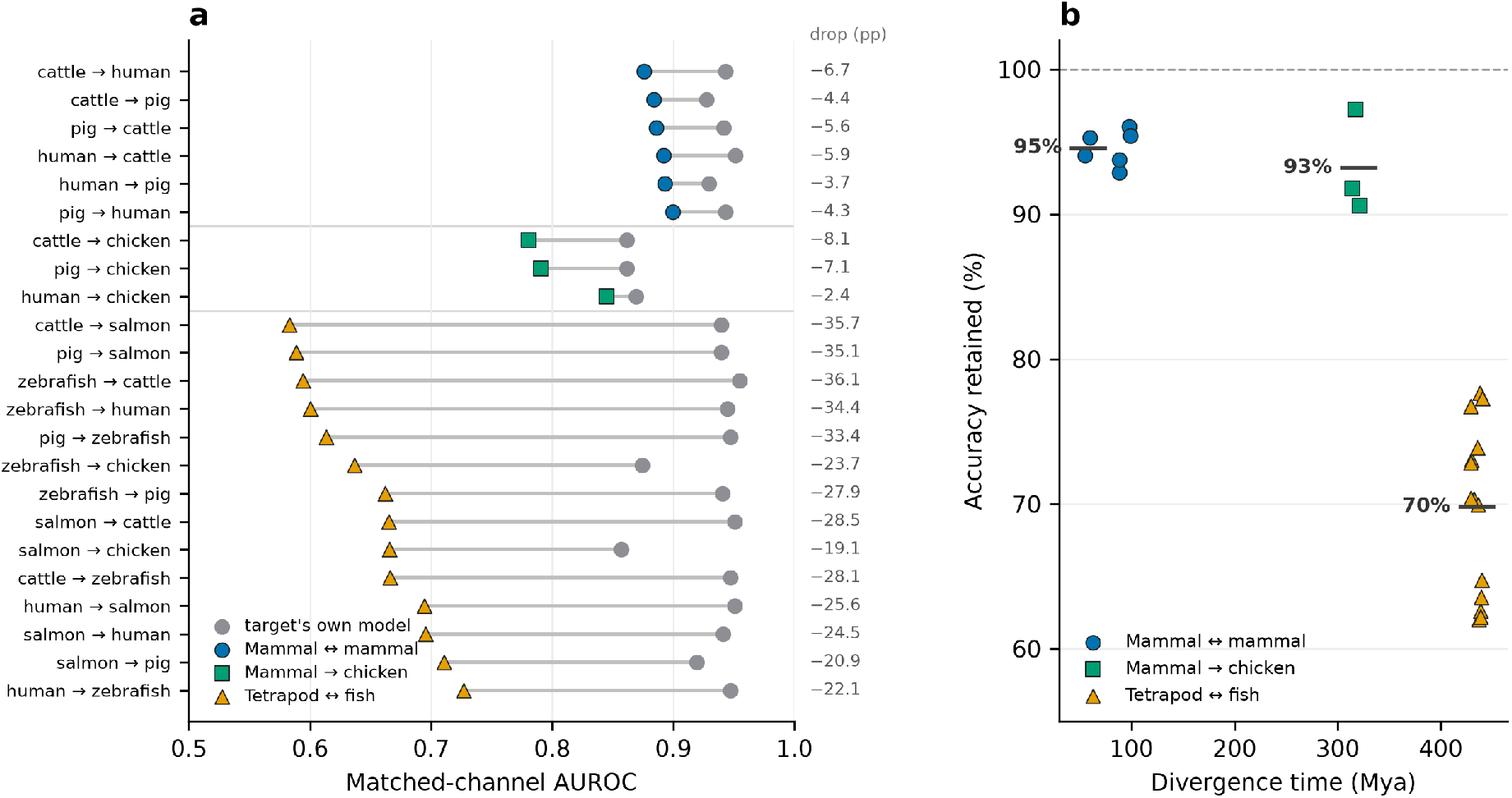
Accuracy given up against the model trained on the target genome. Each transfer is referenced to the target’s own FunFormer-20k on the same channels. **(a)** Per direction, the target’s own AUROC (gray) joined to the transferred AUROC, the drop in pp at the right; directions grouped by bin, ordered by transferred AUROC. **(b)** Retained fraction against divergence time, bars bin means. Transfer retains 95% of the target model’s accuracy among mammals, 93% across the amniote split, but 70% across the tetrapod-fish split. Mammal-to-chicken transfer reaches a low 0.805 mostly because chicken is the hardest genome for any model, its own included (0.865). Only the drop across the tetrapod-fish split is a transfer failure, the targets in that bin scoring 0.932 under their own models (0.945 for the two fish). Color and marker follow the non-mammal partner’s clade.

**Figure S4:**
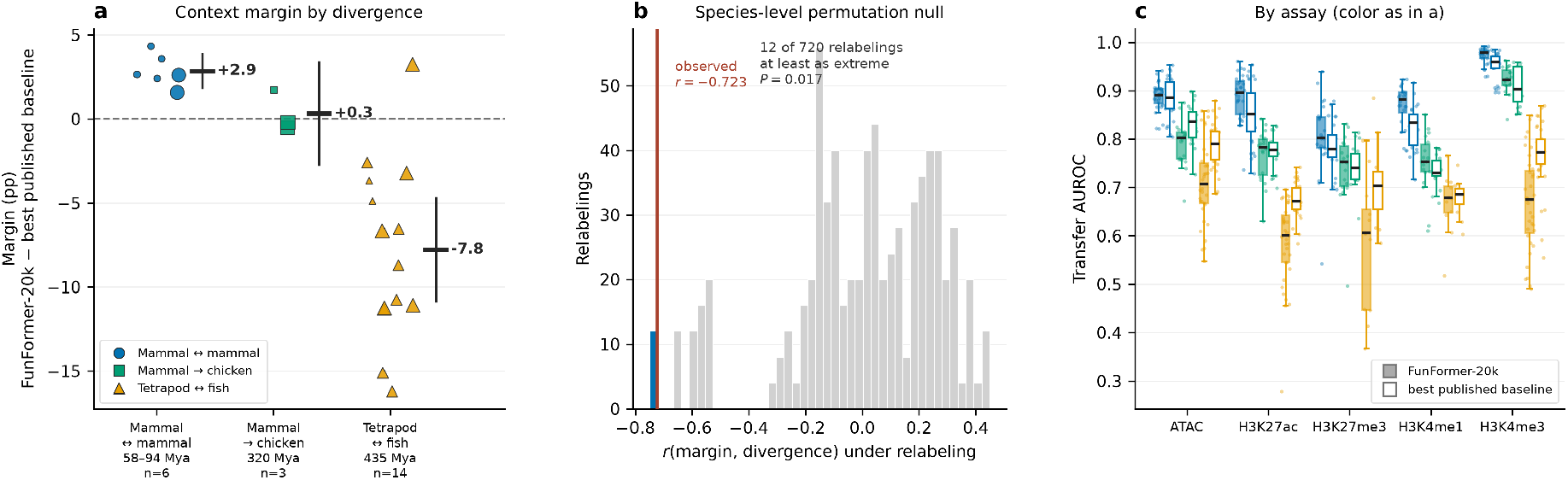
The gradient does not depend on which 1 kb model is the reference. Figure 3a–c with the margin taken against the best published 1 kb baseline instead of FunFormer-1k. Bin means are +2.9, +0.3 and *−*7.8 pp against +3.4, +0.4 and *−*7.8 pp, the pooled correlation *r* = *−*0.723 against *−*0.764, the exact test the same *P* = 0.017 from 12 of 720 relabelings. Sign pattern, reversal point and significance are unchanged. Supplementary Table S3 covers all three references and both assay sets.

**Figure S5:**
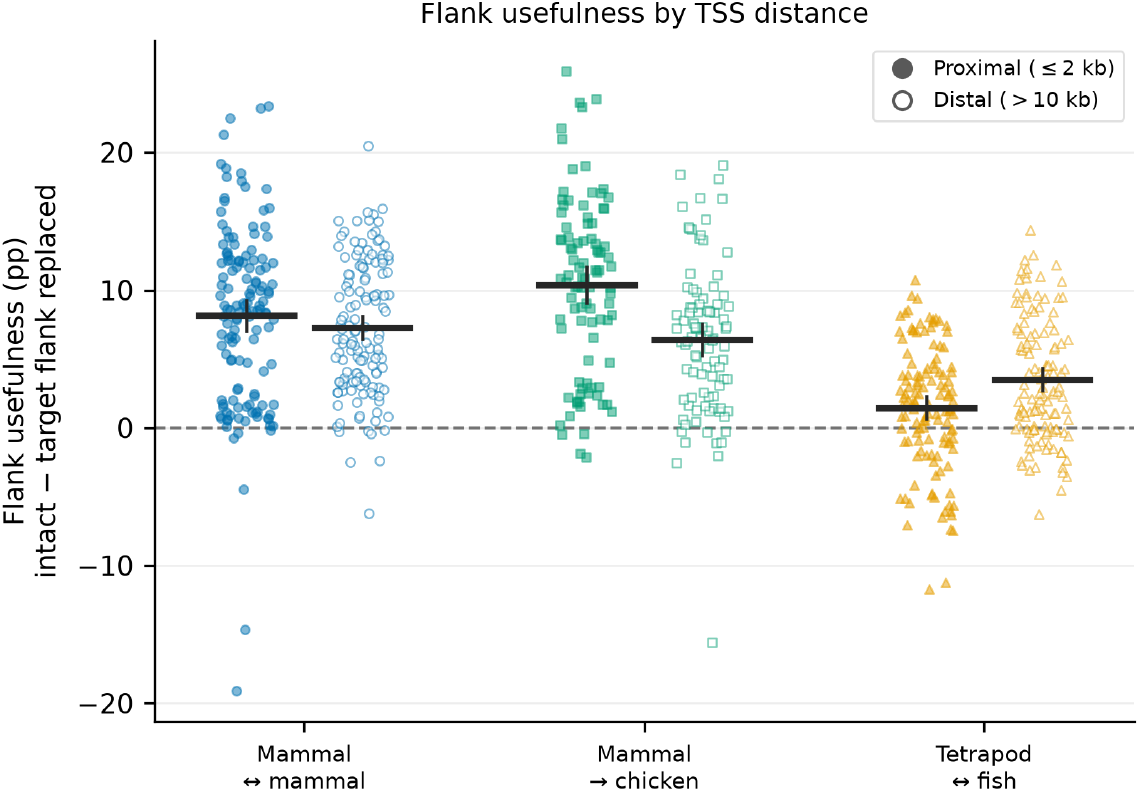
Flank usefulness by distance to a transcription start site. The cost of replacing the target flank with a real, GC-matched sequence from the same genome, per matched channel, split by the strata of Figure 4a. Filled, proximal (*≤* 2 kb); open, distal (*>*10 kb); bar and whisker, the mean and its 95% confidence interval. Among mammals and into chicken the flank is worth more proximally (8.2 against 7.3 pp, and 10.4 against 6.4 pp); at the tetrapod-fish distance the ordering reverses and the proximal flank is worth least of all (1.5 against 3.5 pp).

